# Phase-Separated Droplets Swim to Their Dissolution

**DOI:** 10.1101/2023.07.18.549556

**Authors:** Etienne Jambon-Puillet, Andrea Testa, Charlotta Lorenz, Robert W. Style, Aleksander A. Rebane, Eric R. Dufresne

## Abstract

Biological macromolecules can condense into liquid domains. In cells, these condensates form membraneless organelles that can organize chemical reactions1,2. However, little is known about the physical consequences of chemical activity in and around condensates. Working with model bovine serum albumin (BSA) condensates^3^, we show that droplets swim along chemical gradients. Active BSA droplets loaded with urease swim toward each other. Passive BSA droplets show diverse responses to externally applied gradients of the enzyme’s substrate and products. In all these cases, droplets swim toward solvent conditions that favor their dissolution. We call this behavior *dialytaxis*, and expect it to be generic, as conditions which favor dissolution typically reduce interfacial tension, whose gradients are well-known to drive droplet motion4,5. These results suggest alternative physical mechanisms for active transport in living cells, and may enable the design of fluid micro-robots.

---

Cells control vital bio-chemical processes by creating compartments with distinct compositions. While classical organelles are enclosed by lipid membranes, membraneless organelles hold themselves together^1,2^. The absence of a membrane facilitates their condensation and dissolution^6^, leads to significant interfacial tensions that drive capillary phenomena^7–9^, and enables passive diffusion of solute across their surface^10^. Over the last decade, there has been tremendous progress toward understanding the chemical and physical processes that underlie the condensation of biological macromolecules^11–16^.

Even though biochemical activity is thought to lay at the heart of membraneless organelle physiology, little is known about activity’s impact on condensate stability, properties, and behavior. Experiments in living cells have shown that the fluid properties of some membraneless organelles require ongoing energy consumption^17–19^. Recent *in vitro* experiments have shown that chemical reactions can drive condensation^20^ and suppress Ostwald ripening^21^. Theoretical studies predict that chemical activity can regulate condensation and dissolution^22^, trigger division^23^, and drive chemophoretic motility^24^.

Here, we show that biomolecular condensates can swim toward solvent conditions that favor their dissolution. This process, which we call *dialytaxis*, originates from a generic coupling of phase equilibria and interfacial properties. It emerges naturally whenever droplets, or their surroundings, are chemically active. This simple physical process suggests new mechanisms for active transport within cells, and could enable the design of microscopic fluid robots.

Our model system is schematized in Fig. 1a. We trigger liquid-liquid phase separation by mixing a relatively dilute protein solution (bovine serum albumin (BSA) in PBS buffer (pH 7)) with a concentrated smaller neutral polymer (PEG 4k) (see Materials and Methods). Because this condensation is driven by non-specific depletion interactions^25^, additional proteins, such as enzymes, readily partition into the BSA-rich phase^3^. When the substrate of a partitioned enzyme is available in the dilute phase, these artificial condensates act as chemical micro-reactors. BSA droplets that partition urease catalyze the conversion of urea to ammonia and carbon dioxide: 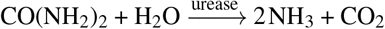 (see Methods and^3,26^).

**FIG. 1.**
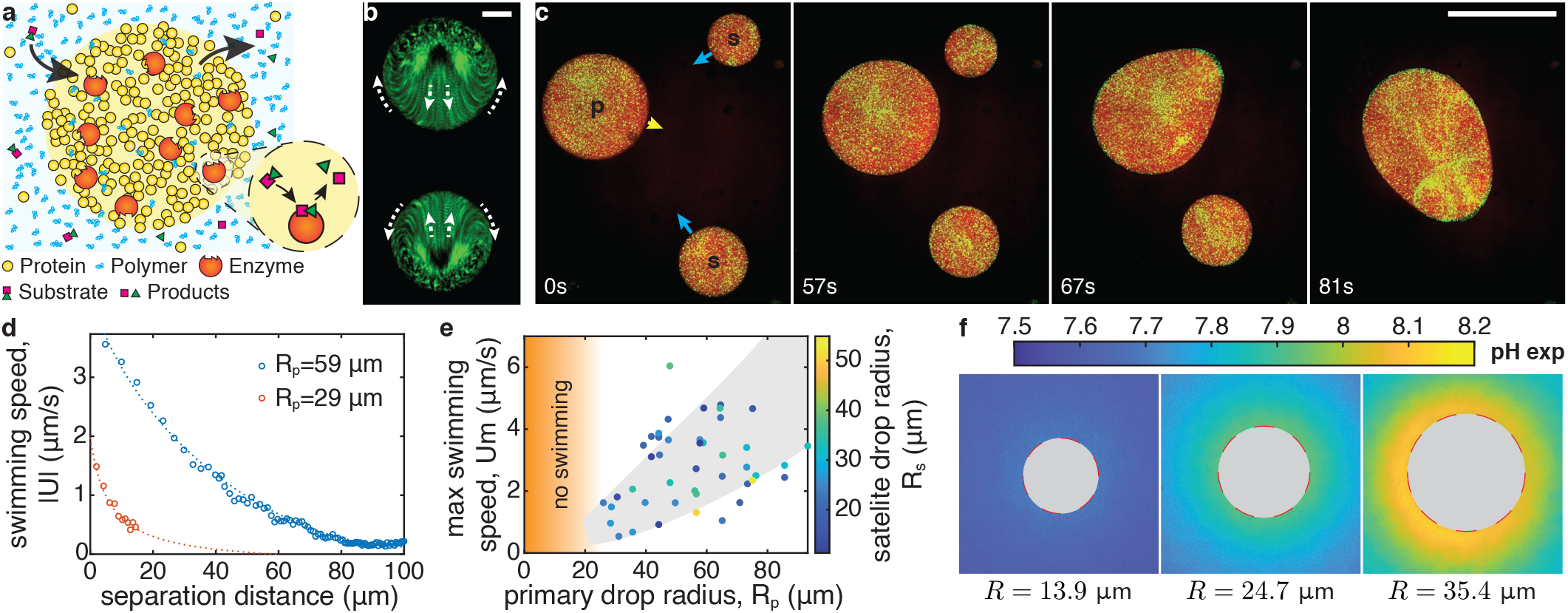
Chemically active protein condensates swim toward each other. **a**. Schematic of a chemically active protein condensate. Polymers act as depletants, triggering condensation. Enzyme-rich droplets act as micro-chemical reactors. **b**. Time projection over 1 min of fluorescent particles inside two adjacent chemically-active protein droplets catalyzing the urea-urease reaction (overall enzyme concentration *ce* = 0.6 μM, substrate concentration *cs* = 100 mM). The droplets are pinned on the surface (PEGDA 700 gel). Arrows indicate the internal flow direction. Scale bar, 10 μm. **c**. Image sequence of chemically active droplets on a non-wetting surface (PEGDA 12k gel, *ce* = 1.2 μM, *cs* = 100 mM, see Movie S1). The primary (p), satellite drops (s) and direction of motion are indicated in the first panel. Scale bar, 100 μm. **d**. Swimming speed of two satellite drops, |*U*|, as a function of the distance from their primary drops of radii *Rp*. Dashed curves are guides to the eye. **e**. Maximum swimming speed, *Um*, as a function the primary drop radius, *Rp*, for 48 swimming-induced coalescence events (*ce* = 1.2 μM, *cs* = 100 mM). The color codes the satellite drop radius, *Rs*, the grey area is a guide to the eye. No swimming was observed for drops below 25 μm in radius as indicated by the orange area. **f**. pH imaging around pinned reacting droplets of increasing size (*ce* = 1.2 μM, *cs* = 100 mM). The fluorescence signal inside the drops is masked by a grey disc, since they do not contain pH dye.

Active urease-loaded droplets develop internal flows that drive collective motility. Attached to a substrate, internal flows are oriented toward neighboring droplets (Fig. 1b)^3^. When the adhesion energy between droplets and substrate is reduced (see^27,28^ and Methods), reacting condensates swim toward each other with speeds on the order of μm/s (Fig. 1c, Movie S1, and^29^).

Like motion driven by attractive potentials (*e*.*g*. gravity or electrostatics), the speed of the droplets depends on their size and separation. Small satellite drops accelerate as they approach a large primary drop (Fig. 1d). Larger primary droplets attract satellites over farther distances. The maximum swimming velocity of satellite drops increases with the radius of the primary droplet (Fig. 1e).

We suspected that these long-range interactions are driven by gradients in the concentration of the chemical species that are consumed and/or produced by the droplet-localized enzymatic reaction. While we could not visualize these small molecules directly, we could observe their downstream effect on the pH. Using the pH-sensitive dye HPTS (see Methods and Extended Fig. 5), we found a cloud of increased pH in steady-state around each reacting condensate, Fig. 1f. Like the motion of satellite droplets, the pH gradient increases with primary droplet size, and is largest near its surface. These size dependent profiles result from the coupled reaction and diffusion of multiple species in and around our droplet micro-reactors as shown in Methods and Extended Fig. 6.

Each of the chemical species involved in the enzymatic reaction could affect droplet motion. To disentangle their contributions, we probed the response of passive (enzyme-free) condensates to controlled gradients in a simple microfluidic device (schematized in Fig. 2a). Enzyme-free condensates were loaded into the central bridge, which was connected to reservoirs of its coexisting dilute phase on one side and dilute phase containing an additional chemical on the other side (see Methods). For ammonia (NH_3_ (aq)), the gradient could be visualized indirectly through its effect on pH, as shown in Extended Fig. 5e-f. Passive droplets in the bridge show an internal flow directed toward the source of NH_3_ (aq) (Fig. 2b), as seen with the active droplets. Suspecting that pH gradients drive motility, we observed the response of passive droplets to sodium hydroxide (NaOH). Again, droplet motion was directed toward increased pH (Fig. 2c). However, this pattern broke down when we explored the impact of the second enzymatic product, CO_2_, which is transformed to bicarbonate in our experimental conditions. In that case, we observed flow directed toward lower pH, away from the slightly basic NaHCO_3_ source at pH7.9 (Fig. 2d). Finally, gradients of the enzymatic substrate, urea, do not drive measureable flow (see Movie S2).

**FIG. 2.**
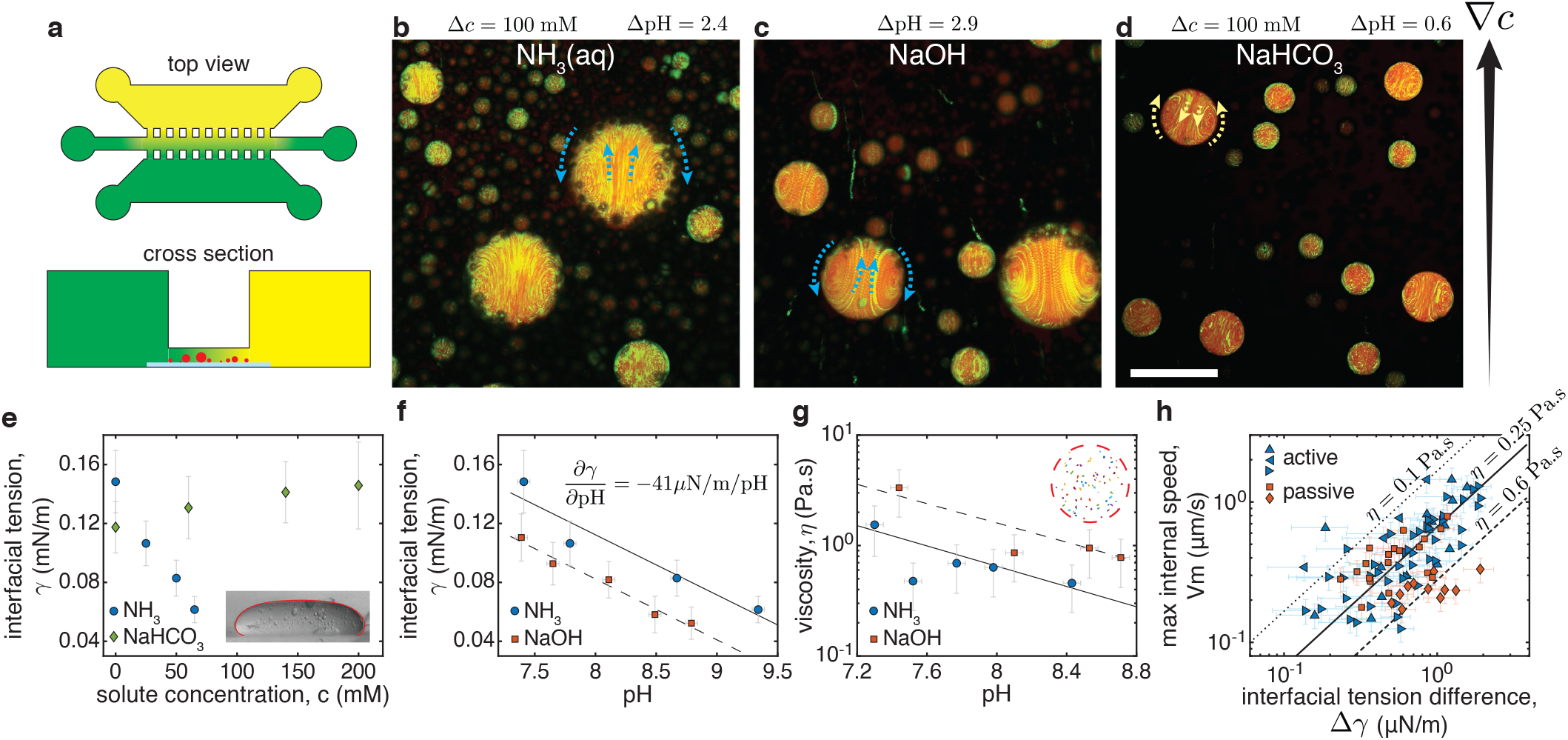
Diverse solute gradients drive droplet motion. **a**. Schematic of the gradient chamber. Bottom reservoir contains the dilute (BSA poor) phase. Narrow bridge is coated with a PEGDA 12k gel and filled with coexisting dilute and dense phases. Top reservoir is filled with dilute phase plus solute. **b.-d**. Time projection over 200 s of fluorescent particles (green) in passive BSA-rich condensates (red) inside the gradient chamber. The extra solutes, the concentration, and pH difference between reservoirs are indicated. The gradient direction is indicated by the black arrow and the internal flow direction by the dashed arrows (see also Movie S2). Drops swim towards the gradient for NH_3_ (aq) and NaOH but away from it for NaHCO_3_. Scale bar, 100 μm. **e**. Equilibrium interfacial tension, *γ*, as a function of solute concentration *c* for NH_3_ (aq) and NaHCO_3_. Measured through the sessile drop method^30^, the inset shows a typical drop flattened by gravity of width 1.8 mm and the Young-Laplace fit. **f**. Equilibrium interfacial tension, *γ*, as a function of pH when adding either NH_3_ (aq) or NaOH. Lines are linear fits. **g**. Droplet viscosity, *η*, as a function of pH for the same solutes, measured with passive microrheology. Lines are exponential fits. The inset show particle tracks longer than 8 sec in a 39 μm drop. **h**. Maximum internal velocity *Vm*, measured as a function of the interfacial tension difference across the drop Δ*γ* = 2*R*(*dγ/d*pH)(*d*pH*/dx*) for 93 individual droplets, both chemically active (blue) and passive ones in a gradient of NH_3_ (aq) (orange). Symbols differentiate independent experiments. The straight lines show the predictions for idealized Marangoni swimmers of viscosities *η* = 0.1, 0.25, 0.6 Pa s. Error bars combine measurement errors and standard deviations (see Methods).

From a fluid mechanics perspective, solute gradients can generate motility through two known mechanisms^31^. They can modify the interfacial tension and drive droplet motion through Marangoni flow^32^. Alternatively, diffusiophoretic motion arises when the interaction of solute and droplet across a diffuse layer generates an apparent slip velocity^33^. While the two mechanisms can work together, Marangoni effects should dominate for micron-scale liquid droplets^4,31^. In that case, droplets swim down interfacial tension gradients and develop internal flows with a maximum speed

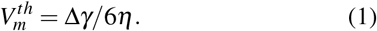

Here, Δ*γ* is the difference in interfacial tension across the droplet and *η* is the viscosity of the more viscous phase^34^. This result neglects the effects of confining surfaces and finite viscosity ratios, and assumes a linear gradient of interfacial tension.

The Marangoni mechanism explains the direction of flow in response to diverse solute gradients. The solutes NH_3_ (aq), NaOH_3_, and NaHCO_3_ each impact the interfacial tension of passive condensates, as shown in Fig. 2e-f (see Methods and Extended Fig. 7). As suggested from Eq. 1 and the observed directions of swimming in Fig. 2, NH_3_ (aq) and NaOH_3_ reduce the interfacial tension, while NaHCO_3_ increases it.

The Marangoni mechanism predicts the observed flow speeds for active (urease-loaded) and passive droplets (in an ammonia gradient). We calculate the interfacial tension differences across droplets by combining pH measurements (see Methods and Extended Fig. 5d) with the measured equilibrium interfacial tensions as function of pH for ammonia (Fig. 2f), Δ*γ* = 2*R*(*dγ/d*pH)(*d*pH*/dx*). We measure the maximum internal flow speed with particle tracking velocimetry (see Methods and Extended Fig. 8). We determine droplet viscosity in passive systems using particle tracking microrheology. Notably, we found droplet viscosities of the order of 1 Pa s that decayed exponentially with pH, by a factor of roughly three per pH unit (Fig. 2g). Combining these results, we plot *V*_*m*_ against Δ*γ* for 93 individual droplets in Fig. 2h. A linear trend is clear for active droplets pooled across all experiments.

For passive droplets, we see a clear trend within individual experiments. Predictions of Eq. 1 are in reasonable agreement with the data for a range of droplet viscosities from 0.1 to 0.6 Pa s. Condensate motility is thus driven by gradients of solutes that affect their interfacial tension. In stark contrast to previous examples of active Marangoni swimmers, however, NH_3_ (aq), NaOH, and NaHCO_3_ are not surfactants^4,5,35^.

The impact of enzymatic products on the interfacial tension emerges from their effect on phase coexistence. An important clue can be found in the behaviour of large active droplets (*R* ≳ 50 μm), which tend to dissolve over long timescales (Movie S3). This suggests that high concentrations of reaction products can dissolve droplets. Indeed, macroscopic experiments with passive PEG-BSA solutions exhibit no phase separation when their pH is increased above about 9, either by the addition of NH_3_ (aq) or NaOH. The coupling between droplet dissolution and motility is vividly demonstrated in Fig. 3a-b and Movie S4. Here, we load one side of a gradient chamber with the dilute phase adjusted to pH 13.8 through the addition of NaOH, and image droplets as the sharp transient pH front moves across the field of view. As the pH front advances, an internal flow oriented toward the basic reservoir accelerates. Eventually, the droplets swim rapidly toward the basic reservoir and dissolve (Movie S4). Together, these observations suggest a direct link between interfacial tension and phase equilibria.

**FIG. 3.**
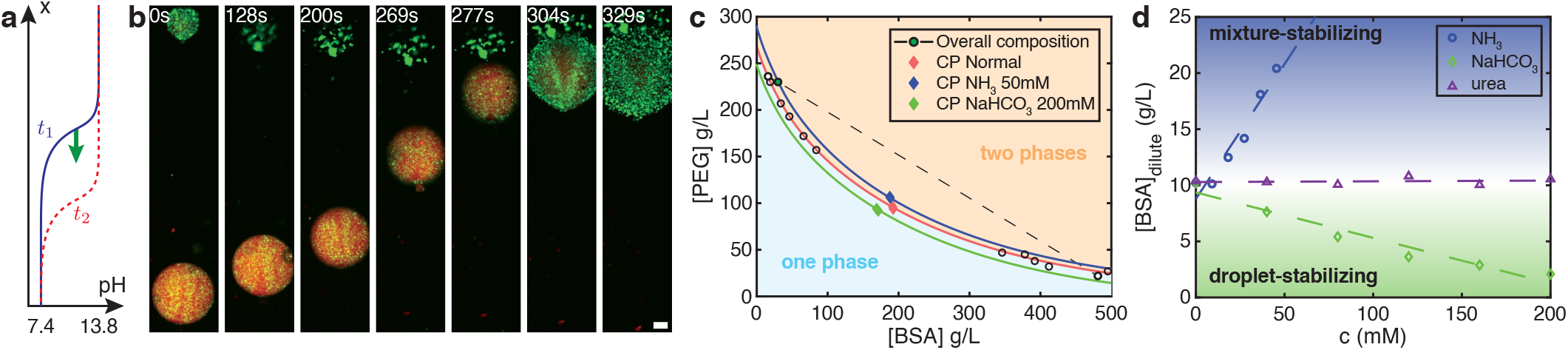
Droplet motility driven by perturbations of phase equilibria. **a**. Schematic of the sharp pH front propagating from top to bottom in b. **b**. Image sequence of passive condensates moving and dissolving in response to the sharp pH front. Scale bar, 10 μm (see also Movie S4). **c**. Phase diagram of the PEG-BSA system. In the absence of solute, circles show the measured binodal^3^ and a red diamond shows the critical point. We prepare our droplets at the overall composition indicated by the green circle, and the system phase-separates along the associated tie line (dashed line and white circles). The background colors and colored curves are guides to the eye. Ezymatic products shift the critical point (blue 50 mM NH_3_ (aq) and green 200 mM NaHCO_3_ diamonds). **d**. Shifts of BSA concentration in the dilute-phase as solutes NH_3_ (aq), NaHCO_3_ and urea are added at fixed overall BSA and PEG concentrations. Dashed lines are linear fits and shaded areas are guides to the eye.

The interfacial tension of coexisting liquid phases depends on their compositions, which can be shifted by solutes. As the overall composition approaches the critical point, the two phases become more similar, and the interfacial tension vanishes^36–38^. The measured phase diagram of our PEG-BSA system is shown in Fig. 3c. Our overall composition (green circle) is close to the PEG-rich arm of the coexistence curve. The system phase separates along the dashed tie-line that connects the compositions of the two coexisting phases. The critical point (red diamond) shifts toward the overall composition with added NH_3_ (aq) (blue diamond), shortening tie-lines, and lowering interfacial tension. NaHCO_3_ shifts the critical point in the opposite direction, stabilizing the droplets and increasing their interfacial tension.

Established theories of polymer mixtures allow us to build a simple model linking interfacial tension and phase equilibria^39^. For segregative phase-separation, we expect *γ*∼(*ϕ*_*m*_/*ϕ*_*c*_ −1)^ν^ where *ϕ*_*c*_ is the total concentration of macro-molecules at the critical point, and *ϕ*_*m*_ is the concentration of macromolecules at the center of the tie-line^38^. Far from the critical point, mean-field theory predicts ν = 3/2. Therefore, given an empirical shift of the critical point with solute concentration, *∂ϕ*_*c*_/*∂c*, we expect a corresponding shift in the interfacial tension, *dγ* ∼ − (*∂ϕ*_*c*_/*∂c*)*dc*. In a solute gradient, the swim velocity will therefore scale like

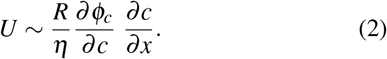

For *∂ϕ*_*c*_/*∂c >* 0, droplets will swim up the solute gradient. Conversely, droplets swim down the solute gradient when *∂ϕ*_*c*_/*∂c <* 0. This simple result captures essential features of our observations. While Eq. 2 was derived for segregative phase separation, we expect shifts of the critical point toward the overall composition to generally reduce interfacial tension. In other words, we expect droplets to swim to their dissolution. We call this process *dialytaxis*, from the Ancient Greek 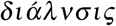 (dialysis), for dissolve, and 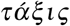 (*taxis*), for arrangement.

While the susceptibility of the critical point to a solute, *∂ϕ*_*c*_/*∂c*, may be challenging to predict from first principles, it can be efficiently measured. For macroscopically phase separated systems, the critical point can be found rapidly using the method of equal volumes (as in Fig. 3, see Methods). However, this is impractical for most biologically relevant condensates, where only small volumes of condensed phases are available. In those cases, the phase behavior can be probed with microfluidics^40^. Alternately, we propose to simply measure dilute phase concentrations of droplet components as a function of solute concentration. Such a measurement can be performed optically (UV absorption or fluorescence) and requires only small amounts of protein. To demonstrate its feasibility, the concentration of BSA in the dilute phase of our system is plotted as a function of ammonia and bicarbonate in Fig. 3d. We clearly see that ammonia drives dissolution while bicarbonate stabilizes droplets, consistent with the observed shifts in the critical point. As a control, we find that urea, whose gradients were not able to drive droplet motility, does not affect the stability of droplets. While a single measurement of the dilute phase concentration has less predictive power than directly measuring the interfacial tension or critical point, it is simple enough to screen large numbers of metabolites for their potential to drive droplet motion.

Dialytaxis provides a simple unifying framework for solutal Marangoni effects in diverse two-phase systems. For example, oil droplets swim toward higher concentrations of ethanol, which promotes dissolution in ternary mixtures of oil, ethanol, and water^41^. Similarly, droplets dispersed in surfactant solutions swim spontaneously^4,42,43^ toward empty micelles that enable their dissolution.

While simple, dialytaxis can drive a rich diversity of behaviors, summarized in Fig. 4. Droplets swim up gradients of solutes that promote mixing, and down gradients of solutes that promote condensation. Active droplets that produce droplet-destabilizing solutes swim toward one another, while active droplets that produce self-stabilizing solutes are expected to swim away from each other. The latter could potentially lead to highly organized steady states. Co-dispersions of active droplets with antagonistic effects on phase equilibria should display non-reciprocal interactions, including predator-prey dynamics^35,44–46^.

**FIG. 4.**
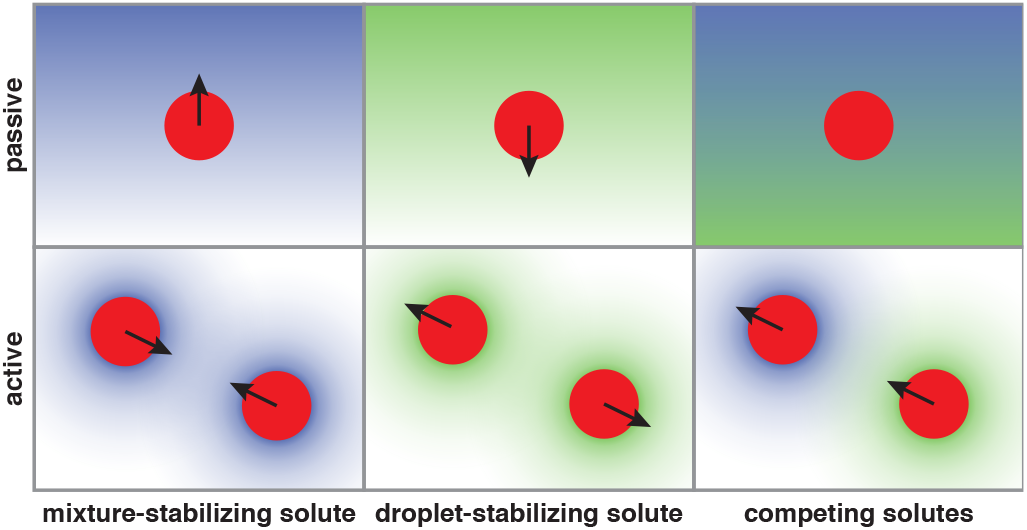
Diverse responses to solubility-perturbing solutes. *(top)* passive droplets are attracted to mixture-stabilizing solutes and repelled from droplet-stabilizing solutes. Opposing gradients can lead to no net motion. *(bottom)* Droplets producing mixture-stabilizing (resp. droplet-stabilizing) solutes swim toward (resp. away from) each other. Droplets producing competing solutes can chase each other.

**FIG. 5.**
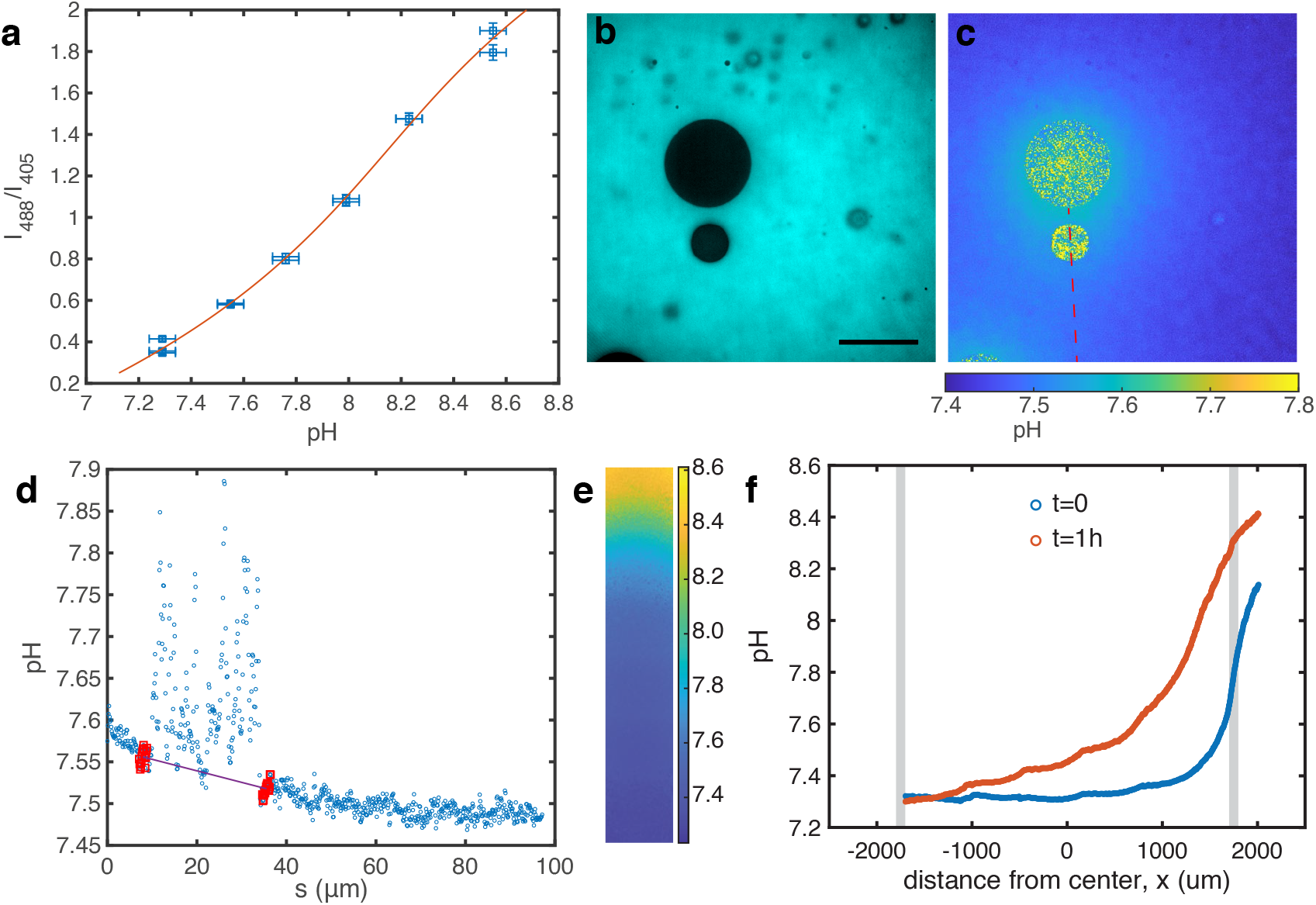
**a**. Calibration curve for the pH measurements using HPTS. The solid curve is a cubic fit of the inverse function pH = f−1(*I*488/*I*405). **b**. Raw fluorescence image after excitation at 405 nm. The dye does not enter the droplets. Scale bar 50 μm. **c**. Same image after conversion of the ratiometric fluorescence values to pH. The pH along the red dashed line is extracted and analyzed to determine the pH gradient around the lower satellite droplet. **d**. pH profile from c. 15 points around each edge of the droplets (red) are used to measure the pH gradient (line). **e**. pH imaging in gradient chamber experiments. The pH in the whole bridge is shown 1 h after the initial injection of dilute phase supplemented with 60 mM of NH_3_ (aq). **f**. pH profile from e averaged along the width immediately after filling the reservoir and 1 h after. Grey bands indicate the edges of the bridge

**FIG. 6.**
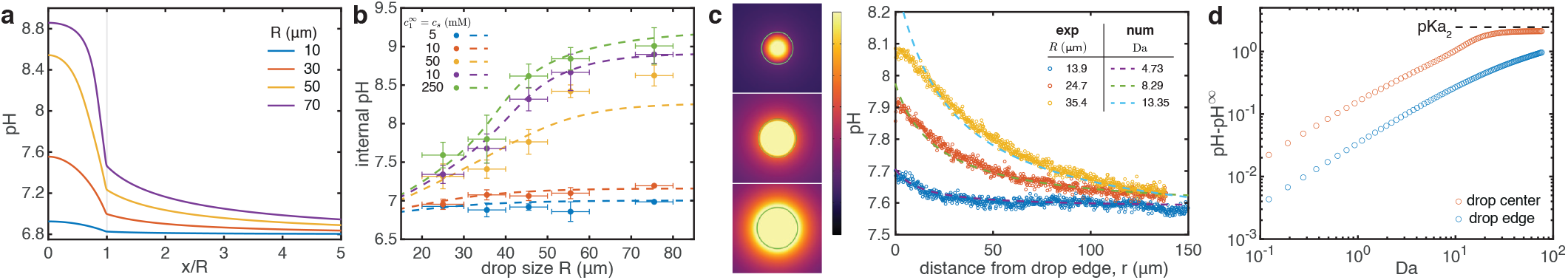
**a**. pH profiles as a function of the dimensionless distance *x/R* as the drop radius *R* increases (pH∞ = 6.8, *η*_*dense*_ = 0.8 Pa s, 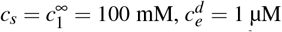, *k*_*cat*_ = 4400 1/s). **b**. Internal pH as a function of the drop radius *R* for various urea concentrations *cs*. Circles are experimental data from^3^, dashed lines are simulations (same parameters as a, except for *cs* that is varied). **c**. Numerical prediction for the pH around the reacting droplets shown in Fig. 1f (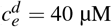, *k*_*cat*_ = 352 1/s, pH∞ = {7.59, 7.59, 7.54} and *η*_*dense*_ = {0.8, 0.3, 0.2} Pa s). Images of the numerical simulations are shown on the left (the green circles indicates the drop edge), and pH profiles outside of the drop extracted from the experiments and simulations are shown on the right. **d**. Increase in pH due to the activity pH pH∞ as a function of the Damköhler numbers Da in the drop center and at the edge (same parameters as a). The black dashed line is pH = pKa_2_.

**FIG. 7.**
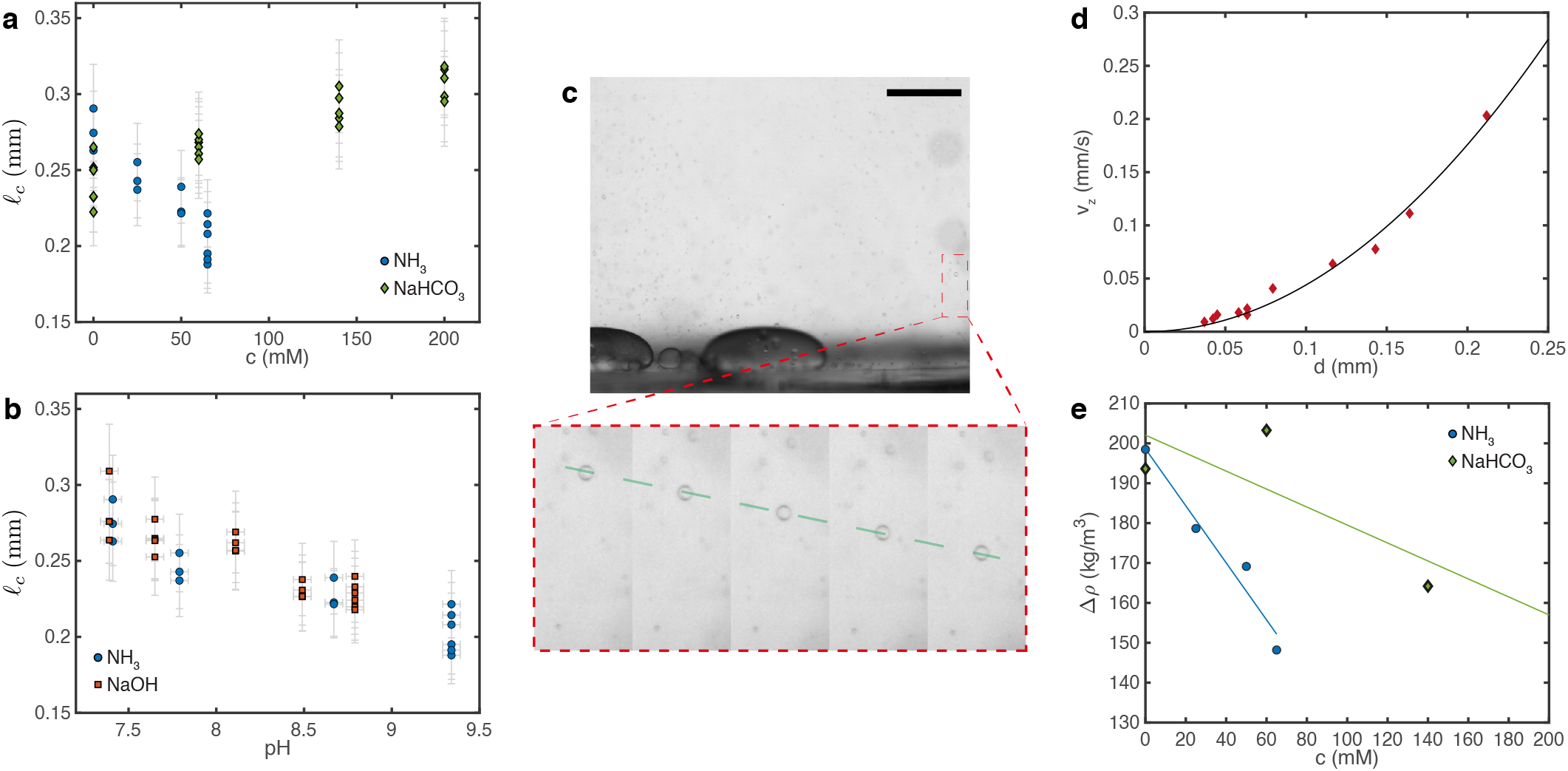
**a-b**. Capillary length *ℓ*_*c*_ as a function of solute concentration *c* and pH for NH_3_ (aq), NaHCO_3_ and NaOH. Each point represents a measurement on one droplet. **c**. Snapshot of a movie looking at the sedimentation of small drops. Scale bar 1 mm. The inset shows an image sequence zoomed on a single drop (*dt* = 7 s). The drop has reached a constant velocity as shown by the dashed line. **d**. Terminal velocity *v*_*z*_ as a function of the drop diameter *d*. The solid curve is a quadratic fit *v*_*z*_ = *Ad*^2^, from which we extract the density difference, Δ*ρ*. **e**. Density difference Δ*ρ* as a function of solute concentration *c* for NH_3_ (aq) and NaHCO_3_. The solid lines are linear fits that we use to compute *γ*, the distance to the fitted lines is our measurement error *δ*(Δ*ρ*).

**FIG. 8.**
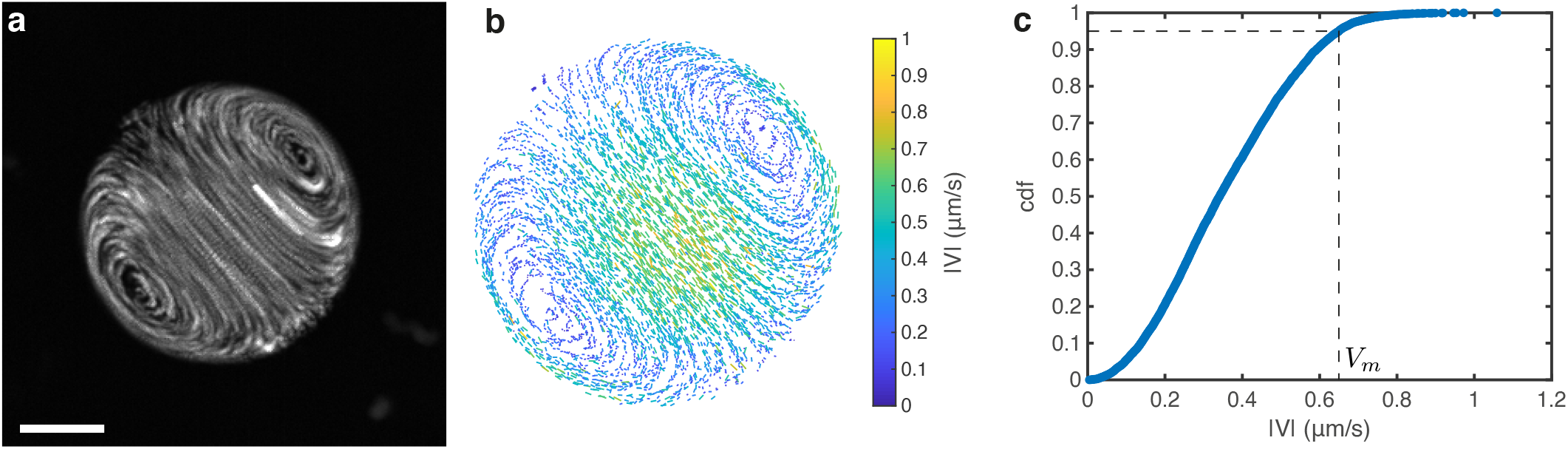
**a**. Time projection of fluorescent particles inside a satellite droplet pinned on a PEGDA 700 coating. Scale bar 10 μm. **b**. Quiver plot after particle tracking for the droplet shown in a. The color codes the magnitude of the internal velocity |*V*|, only half of the tracks are displayed for clarity. **c**. Cumulative distribution function of the internal velocity for the same droplet. Since the highest recorded values are likely to be noise from tracking errors, we define the maximum internal velocity *V*_*m*_ as the velocity larger than 95% of our tracked velocities as shown by the dashed line.

Broadly speaking, dialytaxis suggests new mechanisms of active transport within living cells, independent of molecular motors or the cytoskeleton. We envision multiple contexts where dialytaxis could impact the physiology of membraneless organelles. Most simply, dialytaxis could regulate their coalescence^47^. Further, chemical gradients could contribute to assymmetric inheritence of membraneless organelles during cell division^6^. Similarly, dialytaxis could create symmetry-breaking fluxes of intermediaries for multi-step syntheses, such as ribosome biogenesis^7^. Finally, localized reactions could act as chemical beacons that bring biocondensates together to efficiently deliver diverse reactants.

Moreover, the generality and versatility of this simple chemo-mechanical coupling makes active macromolecular condensates an intriguing candidate platform for micro-robots^48,49^. The results in this manuscript already demonstrate rudimentary capabilities of sensing, actuation, and energy harvesting. Through a complex network of reactions within a single droplet, or through the interactions of diverse droplet types, active droplets should further be able to process information and implement control systems.

## MATERIALS AND METHODS

### Materials

Polyethylene glycol 4000 Da (PEG 4k) (A16151), Polyethylene glycol 12 kDa (PEG 12k) (042635) were purchased from Alpha-Aesar. Bovine serum albumin (BSA) (A7638), Potassium phospate dibasic trihydrate (60349), Potassium chloride (KCl) (60128), 3-(Trimethoxysilyl)propylmethacrylate (440159), 2-Hydroxy-4’-(2-hydroxyyethoxy)-2-methylpropiophenone (410896), Polyethylene glycol diacrylate 700 Da (PEGDA 700) (455008), Jack bean urease (U4002), Rhodamine-B (R6626), 8-Hydroxypyrene-1,3,6-trisulfonic acid trisodium salt (HPTS) (H1529) were purchased from Sigma-Aldrich. Potassium phosphate monobasic (42420) was purchased from Acros Organics. Deuterium oxide (D_2_O) (DE50B) was purchased from Apollo. N,N-dimethylformamide (DMF) (D119), Sodium bicarbonate (S/4240/53), Sodium hydroxide (S/4920) were purchased from Fisher Chemicals. Ammonia 25% (1133.1000), Urea (28876.298) were purchased from VWR. Fluorescent polystyrene particles of diameter 0.2 μm (FCDG003) were purchased from Bangs Laboratories. Polydimethylsiloxane (PDMS) SYLGARD 184 kits were purchased from Dow Corning. All the water used is Milli-Q. Imaging is done on a Nikon Ti2 Eclipse confocal microscope equipped with a Nikon DS-Qi2 camera using 20x air, 40x air, and 60x water objectives.

Polyethylene glycol diacrylate 12000 Da (PEGDA 12k) was synthesized by reacting PEG 12k with acryolyl chloride^50^. Briefly, in a Schlenk flask flushed with nitrogen 10 g of PEG 12k, 68.5 mL of dry dichloromethane, 0.46 mL of triethylamine, and 0.35 mL of acryolyl chloride are left to react overnight while stirring. About half of the content of the flask was then evaporated (Rotovapor R-300 Büchi at 50°C/690 mbar) and the rest was precipitated into ethyl ether. The powder was then vacuum-filtered.

### Condensate preparation

We prepare stock solutions with water: PEG 4k 60%w/v, KCl 4 M, urea 8 M, ammonia 5 M, sodium bicarbonate 1 M, NaOH 3 M, HCl 1 M, potassium phosphate buffer (KP) 0.5 M at pH = 7 (adjusted with pH meter Orion Star A111, Thermo Fisher). BSA and urease stock solutions concentration are measured after their preparation using a UV-vis spectrophotometer (Cary 60 Spectrophotometer, Agilent Technologies). Briefly, different dilutions in water are prepared (typically 200-100-50 for BSA and 15-10 for urease) and their absorbance spectra are measured in a quartz cuvette (UQ-124, Portmann Instruments). Using the extinction coefficients at 280 nm *ε*_*BSA*_ = 43.824 mM^−1^cm^−1^ and *ε*_*urease*_ = 54.165 mM^−1^cm^−1^ we calculate the concentration of each sample and average them to get the stock solution concentration (around 4.5 mM for BSA and 110 μM for urease). PEG 4k, BSA, urea and urease stocks are stored at 4°C.

Condensates are prepared by mixing the appropriate amount of stock solutions and water in a microcentrifuge tube. The total volume for the final solution is typically 1 mL, containing main components at overall concentrations of 23%w/v for PEG 4k, 3%w/v for BSA, 200 mM for KCl and 100 mM for KP (pH 7). The concentrations of the extra components (urease, urea, NH_3_ (aq), NaHCO_3_ and NaOH) are variable as indicated in the main text. When needed, 0.2 μm fluorescent particles at a final dilution of ∼20000 (488 nm excitation, 525 nm emission filter) and Rhodamine-B at 10 μg/mL (561 nm excitation, 600 nm emission filter) are used for visualization. The particles are used for visualizing internal flows and rhodamine is used for visualizing condensates since it strongly partitions inside them. The depletant PEG 4k is always added as the very last component and the solutions are then gently mixed with the micropipette until homogeneous. The final volume fraction of condensates is of the order of *ϕ*_*d*_ = 3*±*2% in the absence of extra components.

### Coatings preparation

The protocols for coating coverslips with PEGDA hydrogels are described in detail elsewhere^27,28^. Briefly, we first silanize the glass coverslips. This is done by washing the coverslips (VWR, #1 thickness) with water, ethanol, and again with water. We then expose them to UV-ozone (ProCleaner, Bioforce Nanosciences) for ∼ 10 min. Then, ∼ 50 μL of 95% v/v ethanol-water solution containing 0.3% v/v of the silane coupling agent 3-(Trimethoxysilyl)propylmethacrylate are spread on the clean coverslips and left to react for 3 min. Finally, the reaction is quenched with ethanol and the coveslips are dried and stored with desiccant.

Subsequently, the silanized coverslip is coated with PEGDA solution containing photoinitiator and cured under UV light. Specifically, we dissolve the photoinitiator 2-Hydroxy-4’-(2-hydroxyyethoxy)-2-methylpropiophenone in water (1%w/v for PEGDA 700 or 4%w/v for PEGDA 12k) by sonicating in a bath sonicator for 30 min at 55°C. The initiator solution is then added to pure PEGDA 700 solution at a 80/20 (v/v) ratio (or to a 40%w/v aqueous solution of PEGDA 12k at a 50/50 (v/v) ratio). We wash a fresh cover slip with water, ethanol, and again with water, and treat it with a commercial hydrophobic coating (Rain-X^®^). About 50 μL of the PEGDA and photoinitiator solution is then added to a silanized coverslip and then covered with a hydrophobic one. The glass-PEGDA 700 sandwich is then cured under 360 nm UV light for 20 min (or 1h for PEGDA 12k) and opened with a scalpel for immediate use, or stored in moist conditions. Our PEG-BSA condensates have a contact angle of ≈ 20° on bare glass, ≈ 100° on PEGDA 700 and ≈ 180° on PEGDA 12k^28^.

### Active condensate experiments

We prepare three different condensate solutions with Rhodamine: solution 1 does not contain any additives, solution 2 contains urease (*c*_*e*_ = 0.6 to 1.2 μM) and fluorescent particles, and solution 3 contains urea (200 mM). Solutions 1 and 3 are then centrifuged until the dense and dilute phases are separated (16000 g, 30 min), and the dilute phases are subsequently extracted and transferred to separate microcentrifuge tubes. In the meantime, solution 2 is kept in a rotator and a PEGDA coated cover slip is prepared. If PEGDA 12k is used, the coating is equilibrated for 30 min with the previously extracted dilute phase of solution 1. The PEGDA-coated coverslip is then inserted into a Chamlide sample chamber (CM-S18-1, CM-S18-4, from Live Cell Instrument) and the chamber is filled with the dilute phase of solution 1 and a small amount of solution 2 (∼ 1 μL). At this point, the sample chamber contains a a very dilute solution of urease-loaded protein condensates without urea. The chemical reaction is triggered by injecting an equal amount of the previously extracted dilute phase of solution 3, which contains urea, and monitored under the confocal microscope.

### Gradient chamber experiments

The gradient chambers are made of PDMS (cured overnight at 40°C with a 10:1 ratio) pressed on a cover slip with a PEGDA 12k coating on the bridge location (see Fig. 2a). The PDMS mold is 3d printed (Pursa SL1) and thoroughly washed and dried before use. The dimensions of the reservoirs are 22×5×1 mm, the bridge is made out of a central stripe of 30×1.5×0.2 mm and connecting teeth of 1.5×1×0.2 mm. The total distance between the reservoirs is thus 3.5 mm. Upon demolding, 2.5 to 3 mm holes are made with a hole puncher to allow fluid injection (circles in Fig. 2a) and the PDMS is gently pressed on a PEGDA coated cover slip to complete the chamber.

As in the active condensate experiment, we prepare three different passive condensate solutions with Rhodamine: solution 1 does not contain any additives, solution 2 contains fluorescent particles, and solution 3 contains the desired chemical (urea, NH_3_ (aq), NaHCO_3_, or NaOH). Solutions 1 and 3 are centrifuged and the dilute phase is extracted and transferred to clean microcentrifuge tubes. Solution 2 is diluted with the dilute phase of solution 1 (usually 1/3 (v/v)) and pipetted into the bridge openings. Capillarity then sucks the fluid in the whole bridge. Solution 1 is then gently pipetted into the first reservoir while capillarity prevents the invasion of liquid in the second reservoir. The chamber is then placed in a custom humidity box filled with a wet sponge and placed under the microscope. Solution 3 is then slowly pipetted into the second reservoir until complete filling. There is some mixing in the bridge at this stage, making the initial conditions unknown, except when pH dye is used. Removing Rhodamine from solution 3 allows us to qualitatively gauge injection-induced mixing. The gradient chamber is left for > 1 h to equilibrate before any measurements are taken (except for the experiment of Fig. 3a-b involving a moving front).

When using a high concentration of NH_3_ (aq) or NaOH, such that pH ≳ 9, solution 3 did not phase-separate and was injected as is. Adding the equivalent amount of NH_3_ or NaOH directly to the dilute phase rather than in the condensate recipe did not yield significant differences, most likely because our overall composition is very close to the dilute phase composition (see Fig. 3c).

### pH measurements

pH measurements are performed either with active condensates or in the gradient chamber with passive ones. The procedure is very similar to what is described above for producing and visualizing droplets, except that instead of Rhodamine, the pH-sensitive dye HPTS is used at 400 μM final concentration. The pH measurment requires a calibration curve that relates the ratiometric fluorescence intensities (see below) to the pH. To this end, six to eight 200 μL calibration solutions are prepared for each experiment by mixing the dilute phases extracted from solution 1 and 3 at equal volumes and altering their pH in the range 7-9 by adding a few μL of NaOH or HCL (measured with Orion Star A111 pH meter). The calibration curve is sensitive to experimental conditions and should therfore be measured under identical conditions as the experiment: in Chamlide CM-S18-4 sample chambers with a PEGDA-coated cover slip, using the same microscope objective (usually 60x water) and performing a z-scan above the coating until fluorescence becomes independant of z.

The pH is determined from the ratios of the fluorescence intensities emitted at 525 nm for two different excitation wavelengths, 405 nm and 488 nm. The laser power and exposure are set to yield a signal to noise ratio above 10 at all pH values while care is taken to minimize sample heating. The raw signals are corrected for vignetting and the sensor noise (intensity with laser off) is subtracted. The calibration is performed at multiple locations in each sample chamber and the whole image is averaged. Extended Fig. 5a shows a typical calibration curve *I*_488_/*I*_405_ = f(pH) whose inverse function pH = f^−1^(*I*_488_/*I*_405_) is fitted by a third order polynomial. This fit is then used to convert the measured signal to pH values in the experiment. As shown in Extended Fig. 5b, HPTS doesn’t enter the droplets, and therefore the ratiometric signal is insufficient to extract meaningful pH values inside droplets. To measure *d*pH/*dx* around satellite drops in active experiments, we extract the pH on a line aligned with the drop swimming direction and fit the pH around the drop (see Extended Fig. 5c-d). For passive gradient chamber experiments, we average the pH on the width of the image (see Extended Fig. 5e-f) and then fit a line around the drop position.

### Reaction diffusion model and numerical simulation

To understand the dependence of concentration profiles on droplet size, we need to account for coupled reaction and diffusion of multiple species in and around our droplet micro-reactors. To that end, we built a multi-component numerical reaction-diffusion model including the reactions of the enzyme with substrate and its product with the buffer^51^.

The system of chemical reactions for our active condensate system is the following (ignoring PO_4_^3–^ and CO_3_^2–^, which are negligible at the experimentally observed pH):

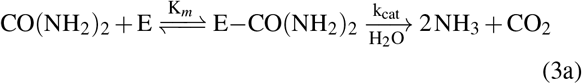

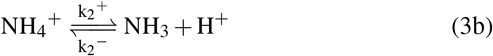

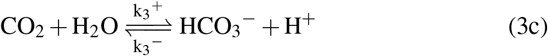

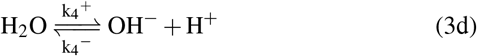

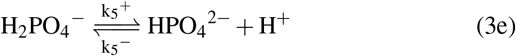

Here a Michaelis–Menten kinetics with parameters *k*_*cat*_ and *K*_*m*_ is assumed for the enzymatic reaction and 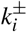 are the reactions rates of the proton exchange reactions.

Assuming no advection (i.e. 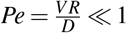1), steady state (i.e. *∂c/∂t* = 0), and denoting *c*_*i*_ the concentration of the compnent *i* with

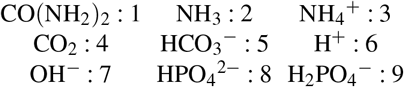

the system of reactive transport equations is:

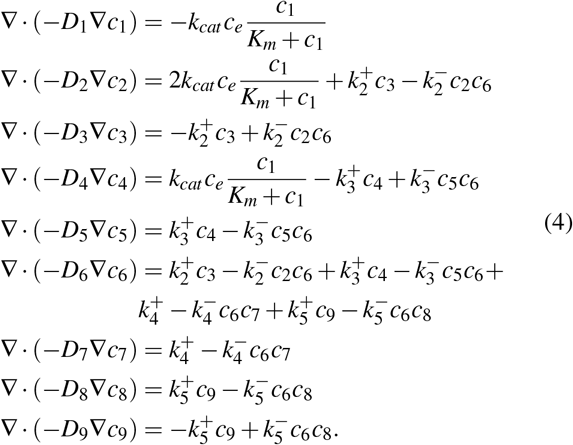

Now following the steps of Ref^51^, we rearrange eq. (4) in order to eliminate the unknown 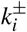:

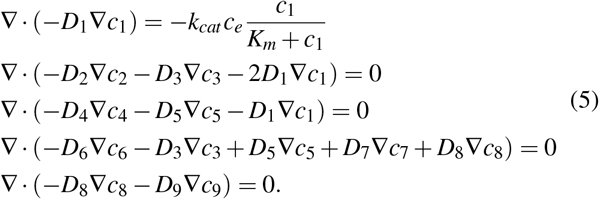

This yields 5 equations with 9 unknowns, and we obtain the 4 missing equations by assuming equilibrium of the proton exchange reaction:

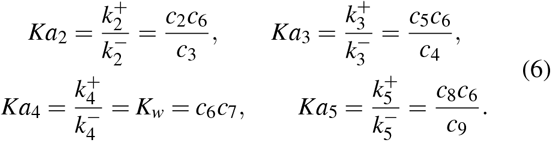

Eqs. (5)(6) are solved numerically for a spherically symmetric droplet (neglecting the surface) with the COMSOL Multiphysics^®^ software, assuming no flux at the origin *∂c*_*i*_/*∂r*|_*r*=0_ = 0 and constant concentrations at infinity 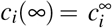 We assume no partitionning for all small molecules (*c*_*i*_(*R*^−^) = *c*_*i*_(*R*^+^)) and infinite partitioning for the enzyme 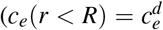 and *c*_*e*_(*r > R*) = 0). This discontinuity is enforced numerically with the smooth step function 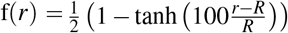 and the mesh is tightened around *r* =*R*. Similarly we expect a discontinuity of all the diffusivities *D*_*i*_ at the condensate edge due to large viscosity contrast between the dense and dilutes phases. Since our diffusing components are much smaller than the macromolecules in both the dense and dilute phases, obstruction theory^53^ rather than the Stokes-Einstein relationship will govern their diffusivity. Following ref^54^, we estimate the diffusivities in the dense and dilute phases using their respective viscosities: 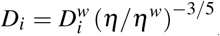 with 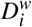the diffusivity of species *i* in water and *ηw* Pa s the viscosity of water. We impose the diffusivity discontinuity numerically using the same smooth step function f.

Extended Table 1 shows the values of parameters kept constant in all the simulations. We further use *η*_*dilute*_ = 24 mPa s for the dilute phase^3^ in all simulations. The concentration of enzymes in the drop is estimated assuming infinite partitioning, i.e. 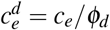. Since the volume fraction of condensate of *ϕ*_*d*_ = 3 *±* 2% can vary substantially there is a large uncertainty on 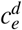. In the far-field the experimental values of urea concentration 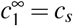 and 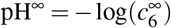 are used while the viscosity inside the droplet *η*_*dense*_ is adjusted within the experimentally measured range 0.1 - 1 Pa s (see values for each simulation below). The catalytic constant *k*_*cat*_ depends on the activity of the urease used (which can decrease as the enzyme ages) as well as the experimental conditions (buffer, pH, temperature). For jack beans urease we expect *k*_*cat*_ ∼ 10^3^ − 10^4^ 1/s^3,55^. Given that the experimental value of both 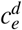 and *k*_*cat*_ are uncertain and that they act together in eq. (5), we use *k*_*cat*_ as fitting parameter.

We first show in Extended Fig. 6a typical numerical pH profiles as we vary the drop size *R*. The pH is larger inside the drop and decreases close to the edge where it exhibits a slope discontinuity; the overall pH magnitude and extent increases with the size of the drop as seen in experiments. We then compare our simulations to experiments. Extended Fig. 6b shows the simulated internal droplet pH and the one measured in^3^. We find a quantitative agreement with a reasonable value of the catalytic activity *k*_*cat*_ = 4400 1/s (all other parameters are known). Finally, we compare our simulation to the data of Fig. 1f for the pH outside of the droplets in Extended Fig. 6c. We find a quantitative agreement for the three drops with a single value of *k*_*cat*_ = 352 1/s. Our model thus captures the radius-dependence of the pH for the three droplet sizes shown in Fig. 1f, {R = 13.9, 24.7, 35.4} μm. The remaining parameters of the model are 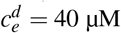, pH^∞^ = {7.59, 7.59, 7.54} and *η*_*dense*_ = {0.8, 0.3, 0.2} Pa s. The *k*_*cat*_ is lower than expected, which suggests that our enzyme batch may have lost some of its activity.

To understand this radius-dependence, we come back to eq. (5). Since the pH increase is driven by the enzymatic reaction inside the droplet (where *D*_*i*_ are constants), we isolate it and make it dimensionless using 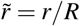 and 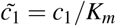:

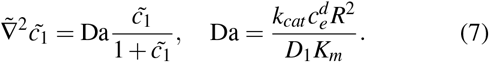

This equation is governed by a single dimensionless number, the Damköhler number Da, which compares the diffusion and reaction timescales. If Da ≪ 1, diffusion is much faster than the reaction and the latter becomes unnoticeable (no pH increase). The reaction speed becomes significant only when Da ≳ 1. Since Da ∼ *R*^2^, the drop size has a strong influence on the transition between the diffusion- and reaction-dominated regimes. Extended Fig. 6d shows the simulated pH increase as a function of Da inside and at the edge of the drop. The pH increases noticeably (by more than 0.1) only for Da > 1. The pH increase eventually stalls when the system reaches the pKa of NH_3_, which effectively becomes the dominant buffer of the system, a feature also present in the experiment (Extended Fig. 6b). Finally, while for Da ≪ 1 the substrate has time to diffuse inside the droplet before being consumed, for Da ≫ 1, the reaction rate is so fast that it is consumed as soon as it reaches the drop edge. We therefore expect the total rate of products released to scale with the volume of the condensate in the first case but with the surface area of the condensate in the latter.

**TABLE I.**
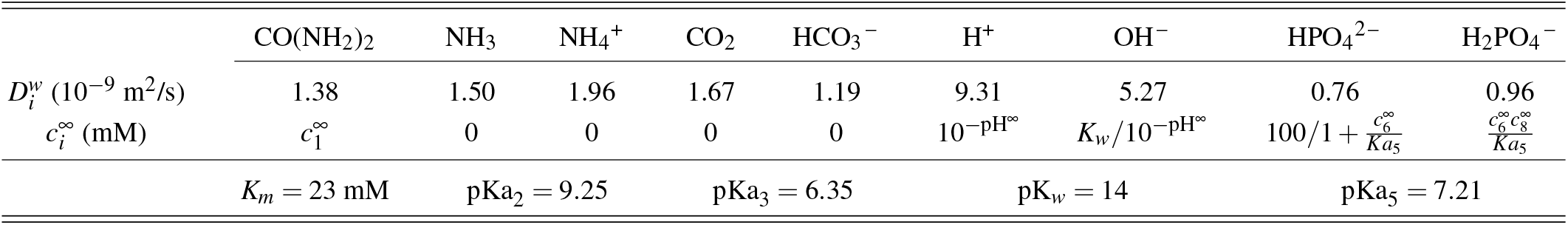
Parameters for the simulation of eqs. (5)(6). Diffusion coefficient and pK are extracted from^52^, urea-urease Michaelis-Menten constant from^3^.

### Interfacial tension, viscosity and internal velocity measurement

#### Interfacial tension

We use the sessile drop method to measure the interfacial tension between the dilute and dense phases^3,30^. We prepare large samples (∼ 2 mL) of passive PEG-BSA condensates at various concentrations of NH_3_ (aq), NaHCO_3_ and NaOH. We then centrifuge each sample, separate the dense and dilute phases, and measure the dilute phase pH. In the meantime, we cut a glass cover slip into 8×8 mm pieces and coat them with PEGDA 12k (see procedure above). The coated glass is then inserted into a standard quartz cuvette that is then filled with dilute phase. We then pipette ≈ 0.5 −1 μL of dense phase at multiple locations in the cuvette to form large condensates as shown in the inset of Fig. 2e. These drops are large enough to be flattened by gravity and we can measure their capillary length 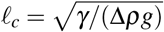 by fitting their shape to the Young-Laplace equation. The experimental setup and fitting code are described in^30^.

The capillary lengths of individual drops are shown in Extended Fig. 7a-b. To calculate the interfacial tension 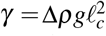 we then measure the density difference between the dense and dilute phases, Δ*ρ*, through the sedimentation velocity of drops of various sizes. We generate clouds of small droplets by mixing the content of the cuvette with a pipette. After some time the background flow vanishes and we record the sedimentation velocity *v*_*z*_ of droplets of different sizes *d* as shown in Extended Fig. 7c. For isolated drops, assuming that they have reached their terminal velocity and that *η*_*dense*_ ≫ *η*_*dilute*_, we extract the density difference from a quadratic fit of the data: *v*_*z*_ = Δ*ρgd*^2^/(18*η*_*dilute*_) (see Extended Fig. 7d). We plot the extracted density difference in Extended Fig. 7e. Given the large variability, we fit the measured Δ*ρ* with a line and use the fit value to calculate *γ* in Fig. 2e-f, where we used the value of *ℓ*_*c*_ obtained by averaging over individual droplets. The vertical error bars in Fig. 2e-f combine measurement errors on both Δ*ρ* and *ℓ*_*c*_, as well as the standard deviation on *σ*_*ℓ*_*c* for the different droplets:

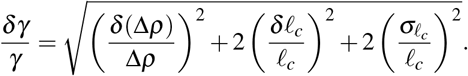

The pH meter measurement error *δ*(pH) = 0.05 is used as the horizontal error bar in Fig. 2f-g.

#### Viscosity

We use passive microrheology to measure the viscosity in the dense phase. For each concentration of NH_3_ (aq) and NaOH, we deposit a diluted passive condensate solution in a sealed sample chamber. Specifically, for each concentration we prepare two solutions of passive PEG-BSA condensate: solution 1 contains fluorescent particles (diameter 0.2 μm) while solution 2 doesn’t. We centrifuge solution 2 and separate the two phases. In the meantime, we prepare a cover slip with a PEGDA 700 coating. We make several sample chambers by pressing an Invitrogen™ Press-to-Seal™ silicone isolator on the coated cover slip. For each concentration, we deposit ∼ 20 μL of the dilute phase extracted from solution 2 in a chamber and then a small amount of solution 1 (∼ 0.2 μL). We then seal the chambers by pressing a clean cover slip on top. The sealing must be excellent to avoid evaporation, which drives flows that hinder the particle diffusivity measurement.

The motion of the particles in a 2d plane is then recorded under the microscope with a 60x water objective. The recording framerate was adjusted between 5 fps and 0.5 fps depending on the sample viscosity. Particle tracking is done with a custom Matlab code available at https://doi.org/10.5281/zenodo.4884194. The particle diffusivity is extracted from the tracks using the optimal estimator from Ref^56^ corrected for a constant background flow:

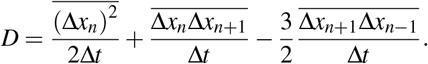

Here,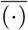 denotes an average over all tracks (all particles, all time points, both x and y direction), Δ*x*_*n*_ the displacements of a tracked particle between frames *n* + 1 and *n* and Δ*t* the time between two frames. We further checked that the estimated diffusivity agreed with the mean squared displacement after correcting the tracks for the average background flow of the experiment, if present. The viscosity is then calculated from the Stokes-Einstein relationship *η* = *kT/*(6*πaD*) with *k* the Boltzmann constant, *T* the absolute temperature and *a* = 0.1 μm the particle radius. This measurement is performed for several individual droplets in a given sample. The measurement reported in Fig. 2g is the average value over all droplets (*n >* 6) and the vertical error bars combine measurement errors on *D* and the standard deviation *σ*_*D*_ for the different droplets 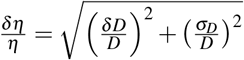

#### Internal velocity

The internal velocity of drops pinned on the surface is measured through particle tracking velocimetry in experiments where pH gradients are recorded (using the 60x water objective, see Extended Fig. 8a). The tracking is done in a 2d plane close to the drops center with the same in-house MATLAB^®^ code as the microrheology experiments.

By differentiating the tracks we get the instantaneous velocity of each tracked particle as shown in Extended Fig. 8b for a satellite drop in an active experiment. We recover the characteristic internal flow of Marangoni swimmers with its pair of vortices. Theory for Marangoni swimmers in infinite space subject to a linear interfacial tension gradient yields simple predictions for the swimming velocity and maximum internal velocity (reached at the drop center and edge, as shown in Extended Fig. 8b)^34^. Since we use pinned drops for pH measurements (that do not swim), we compare the maximum internal velocity to the theoretical prediction. To measure the maximum velocity in the presence of some tracking error (the fastest recorded velocities are likely to be noise), we compute the cumulative distribution function of the velocity magnitude |*V*| shown in Extended Fig. 8c. We define the maximum internal velocity |*V*_*m*_| as the velocity greater than 95% of all the recorded velocities, *i*.*e*. the velocity for which the cumulative distribution function is 0.95. The vertical error bar in Fig. 2g is a measurement error (set to 20%), while the horizontal error bar is a combination of measurement errors on *dγ/d*pH and *d*pH/*dx*.

### Critical point measurement

To measure the critical point of our BSA-PEG system, we first prepared a suspension with the highest possible volume fraction of droplets given our stock solutions. Specifically, we used 17%w/v of PEG 4k, 13%w/v of BSA, 200 mM of KCl, 100 mM of KP (pH 7) to make the droplets. And to see the effect of additional compounds, either 50 mM of NH_3_ (aq) or 200 mM of NaHCO_3_ were added. The suspension was then centrifuged and equal volumes of dense phase and dilute phase were transferred to an empty microcentrifuge tube, resulting in 50% volume fraction of droplets and placing us at the center of the tie-line. The resulting solution was then diluted with buffer containing just 200 mM of KCl and 100 mM of KP (pH 7) and the additional components (50 mM NH_3_ (aq) or 200 mM NaHCO_3_), if any. Upon dilution, the droplet suspension was gently mixed and centrifuged to see whether the solution was still phase-separated. If so, we checked whether the volume fraction of dense phase remained at 50% and if not, the volumes of the two phases were adjusted to achieve 50% volume fraction. This dilution process was repeated in small increments until the solution no longer phase separated. At that point, the composition of this final solution was analyzed by measuring the concentration of BSA with UV-vis spectroscopy (see above) and the concentration of PEG 4k with NMR spectroscopy^3,57^. Briefly, 50 μL of the solution and 30 μL of DMF were dispersed in 570 μL of D_2_O and placed in Schott NMR sample tubes. ^1^H NMR spectra were acquired using a 300MHz instrument (300 Ultrashield, Bruker) and the resulting data were analyzed using the MestreNova software (Mestrelab Research). The PEG 4k concentration was determined by comparing the integrals of the two DMF methyl peaks around 3 ppm (6 protons per molecule) to the intensity of the PEG peak around 3.7 ppm (∼ 354 protons per molecule).

## ACKNOWLEDGMENTS

We thank Magdalini Polymenidou for suggesting the term *dialytaxis*, Iacopo Mattich for help with 3d printing, and Gianna Wolfisberg for help with NMR measurements. This work was supported by grants from the Swiss National Science Foundation (National Center of Competence in Research Bio-Inspired Materials to ERD, grant number 172824 to ERD, and grant number 202214 to AAR).

## AUTHOR CONTRIBUTIONS STATEMENT

EJP, AT, and CL performed the experiments. EJP, RWS, AAR, and ERD analyzed and interpreted the experiments. EJP developed the reaction-diffusion model and simulations. ERD conceived and supervised the project. EJP and ERD wrote the paper with inputs from all authors.

